# Volumetric montaging of optical coherence tomography in human retinas

**DOI:** 10.64898/2026.02.21.706699

**Authors:** Raymond Fang, Fengyuanshan Xu, Daniel Kim, Ronald Zambrano, Alexander Lam, Raphael Tinio, Cheng Sun, Christopher K. S. Leung, Joel Schuman, Rukhsana G. Mirza, Hao F. Zhang

**Affiliations:** Department of Biomedical Engineering, Northwestern University, Evanston IL; Glaucoma Service, Wills Eye Hospital, Philadelphia, PA; Department of Ophthalmology, Hong Kong University Eye Centre, Hong Kong, People’s Republic of China; Department of Ophthalmology, Northwestern University Feinberg, Chicago IL; Department of Mechanical Engineering, Northwestern University, Evanston IL; Sidney Kimmel Medical College, Thomas Jefferson University, Philadelphia, PA; School of Biomedical Engineering, Science and Health Systems, Drexel University, Philadelphia PA

**Keywords:** Data processing, optical coherence tomography, retina, three-dimensional, volumetric montaging

## Abstract

Optical coherence tomography (OCT) has transformed clinical eye care by providing high-resolution volumetric imaging of the retina. Recently, ultrawide-field-of-view (FOV) OCT played an increasingly significant clinical role; however, most clinical OCT systems offer only a rather limited FOV. We increased the FOV of clinical OCT by volumetrically montaging multiple OCT datasets in three dimensions (3D). We performed volumetric montaging by representing the internal limiting membrane (ILM) and retinal pigment epithelium (RPE) in each volume as point clouds and using these point clouds to compute transformations that map each volume to a common reference frame. We validated our methodology using datasets from three institutions with different OCT hardware and data-acquisition procedures. Using the mean surface distance between point clouds, we found the error in montaging was less than the lateral pixel size. Our method enabled existing clinical OCT to achieve ultrawide FOV imaging without any hardware modification.

## I. Introduction

Optical coherence tomography (OCT) is a non-invasive imaging modality that has revolutionized the management of nearly all ocular diseases [1-3]. OCT detects the interferogram from short-coherence-length light to achieve depth-resolved information with microscopic axial resolution. Transversely scanning the beam in the lateral dimension further generates three-dimensional (3D) volumetric images. The field-of-view (FOV) is determined by the lateral scanning size, and the imaging depth is determined by the spectrometer design or the swept-source laser used [4, 5]. Although the FOV can be expanded by mechanically scanning a larger area, this comes at the expense of sparser sampling [6-8] and fringe washout due to fast mechanical scanning [9]. In highly curved eyes with long axial eye length, OCT imaging depth is often unable to capture the full FOV without mirror artifacts [10], [11].

Because many retinal diseases manifest in the peripheral retina, it is clinically important to expand OCT’s FOV. Pathological myopia is a condition in which an increased FOV is beneficial, helpful in locating posterior staphylomas, characterizing abnormal posterior scleral curvature, dome-shaped macula, identifying retinoschisis, retinal breaks, as well as potentially detecting retinal detachment [12-15]. Additionally, expanding the FOV to include the peripheral retina improves the management of diabetic retinopathy and vascular diseases. Recent studies suggest that peripheral retinal nonperfusion helps grade diabetic retinopathy [16, 17]. Consequently, much research has focused on increasing OCT’s FOV.

One method to increase the FOV is to redesign OCT’s optical and hardware components. Researchers have reported using custom-designed widefield lenses to increase the FOV to over 90 degrees [18, 19]. Additionally, robotic OCT has been used to increase the FOV and to locate peripheral regions of interest [20-24]. Unfortunately, many of these customized OCTs are not yet commercially available or are prohibitively expensive. The vast majority of clinical OCTs still have limited FOVs.

Another method to increase the FOV is to montage multiple overlapping volumetric datasets in existing OCTs. The primary advantage of montage-based designs over hardware-based designs for increasing the FOV is the higher scan density. To image a wider FOV in a short time, which is necessary for patient comfort, the lateral sampling density of widefield OCT becomes increasingly sparse. Less-dense sampling results in reduced signal quality and underutilization of OCT’s high resolution. Without sufficiently dense scanning, OCT may fail to visualize smaller retinal features adequately [6-8].

The primary drawback of montage-based methods is the artifacts from the distortion caused by projecting spherical retinal anatomy onto planar projections from multiple OCT volumes [25, 26]. Despite this drawback, montaging remains an attractive way to increase the FOV of existing clinical OCTs.

Several investigators have applied montaging to viewing retinal projections across a wider region of the eye. Most studies on increasing the FOV focused on stitching two-dimensional (2D) *en-face* projection images of the retina [27, 28]. Other investigators have attempted to montage 3D volumes; however, these studies were often performed *ex vivo* and only accounted for coordinate translations between volumes [29-31]. Due to the nature of eye motions between OCT scans, rotation between volumes must be accounted for. Mori *et al*. used montaging to visualize retinal cross-sections over a wider FOV. However, they only merged individual B-scans rather than the full volume [32]. Ji *et al*. developed a method for 3D montaging volumetric images of the skin by flattening each volume and determining the translation between the flattened volumes [33]. However, flattening the volumes distorts the 3D topology of the structure, which is particularly important for understanding the retina’s curvature. So far, there is a lack of a method for 3D volumetric montaging of clinical retinal OCTs.

In this study, we volumetrically montaged up to 36 OCT volumes in 3D to increase the FOV of clinical OCTs. For 3D volumetric registration, we represented each volume as a point cloud. We determined the rigid transformations and displacement vector fields needed to map all volumes to a common reference frame. Compared with previous attempts to combine OCT volumes, this study factors in rotations between acquired volumes. We demonstrate the effectiveness of our method using datasets acquired from multiple institutions, with varying data-acquisition patterns and OCT devices.

## II. Methods

### A. Data acquisition

We acquired data from Hong Kong University (HKU) Eye Centre, Wills Eye Hospital, and Northwestern Memorial Hospital. All studies conducted at these institutions were in accordance with the Declaration of Helsinki and included written informed consent. At HKU Eye Centre, we imaged three eyes with high myopia and one normal eye from four subjects, with approval from the Hong Kong Hospital Authority Research Ethics Committees. Nine 3D OCT volumes were acquired from each eye with a spectral-domain OCT (Spectralis2, Heidelberg Engineering, Germany). Each volume was acquired with a 30° × 28° FOV, which is approximately 10 mm × 10 mm in the eye. Each volume consisted of 135 B-scans, each with 496 A-lines. After acquiring the fovea-centered scan, the center of focus was translated by ±10° in the horizontal direction to acquire two more volumes. From these three positions, the center of focus was further translated by ±10° in the vertical direction for 6 additional volumes.

At Wills Eye Hospital, we imaged eight eyes from seven subjects with approval from the Wills Eye Hospital Institutional Review Board (IRB) and Ethics Committee. Seven healthy subjects with refractive errors ranging from 0 to -6 diopters underwent spectral-domain OCT imaging (Cirrus HD-OCT, Zeiss) using the macular cube and optic disc cube scanning protocols. Each volume consisted of a 6.0 mm × 6.0 mm FOV (approximately 20° × 20°), with 200 B-scans and 200 A-lines per B-scan. We acquired up to 36 volumes with approximately 20% adjacent overlap per volume, covering FOVs of up to 100 degrees per eye. For the 36-volume acquisition, 18 macular cube and 18 optic disc cube volumes were acquired.

At Northwestern Memorial Hospital, we imaged 26 eyes from 13 subjects with approval from the Northwestern IRB. We imaged both eyes of each subject using spectral-domain OCTA (Maestro2, Topcon) with a 6.0 mm × 6.0 mm FOV, 360 B-scans, and 360 A-lines per B-scan. We first obtained a macula-centered scan using internal fixation targets. We instructed subjects to look slightly off-center at the internal fixation target until only 30% of the first macula scan was included. We acquire volumes superior, inferior, temporal, superotemporal, and inferotemporal to the macula. This procedure was repeated with an optic nerve-centered scan with additional volumes in the nasal, superonasal, and inferonasal directions, for 9 scans.

### B. Volumetric montaging overview

After acquiring multiple OCT volumes with partially overlapping regions, we merged the 3D structural data volumetrically. The challenge of montaging multiple datasets reduces to determining the relationship between the local reference coordinates of each volume and a common global coordinate, which is the coordinate system of the first OCT volume (usually the centermost volume).

For the local reference coordinate of each OCT volume *V*_*n*_, we defined the *x*-axis as the fast-scanning axis, the *y*-axis as the slow-scanning axis, and the *z*-axis as the axial axis. Given two adjacent 3D dataset volumes, *V*_*s*_ and *V*_*m*_, we first determined their relative positions and orientations. After that, we transformed the coordinate of *V*_*m*_ (moving volume) to the coordinate of *V*_*s*_ (stationary volume).

Figure 1 shows the flowchart for montaging partially overlapping volumes *V*_*s*_ and *V*_*m*_. In **Step 1**, we segmented the internal limiting membrane (ILM) and retinal pigment epithelium (RPE) for each volume. In **Step 2**, we represented the 3D coordinates of the ILM and RPE segmentation as a point cloud. In **Step 3**, we identified common landmark points in the *en-face* projection of the two volumes. In **Step 4**, we used the 3D coordinates of the landmark points to estimate an initial rigid transformation that aligns *V*_*m*_ to *V*_*s*_. In **Step 5**, we identified the overlapping region between *V*_*s*_ and *V*_*m*_ sharing the same *x* and *y* coordinates in the *V*_*s*_ coordinate system. We registered the point clouds of *V*_*s*_ and *V*_*m*_ in the overlapping region to refine the rigid registration generated in **Step 4**. In **Step 6**, we divided the *V*_*m*_ point cloud into strips. We optimized the transformation of each strip to the *V*_*s*_ point cloud to generate a displacement vector field used for elastic registration of *V*_*m*_ to *V*_*s*_. In **Step 7**, we applied the rigid transformation and displacement vector field to the OCT volume corresponding to *V*_*m*_ to generate a composite merged OCT volume with *V*_*s*_. More details for each step are described in the following sections.

**Fig. 1.**
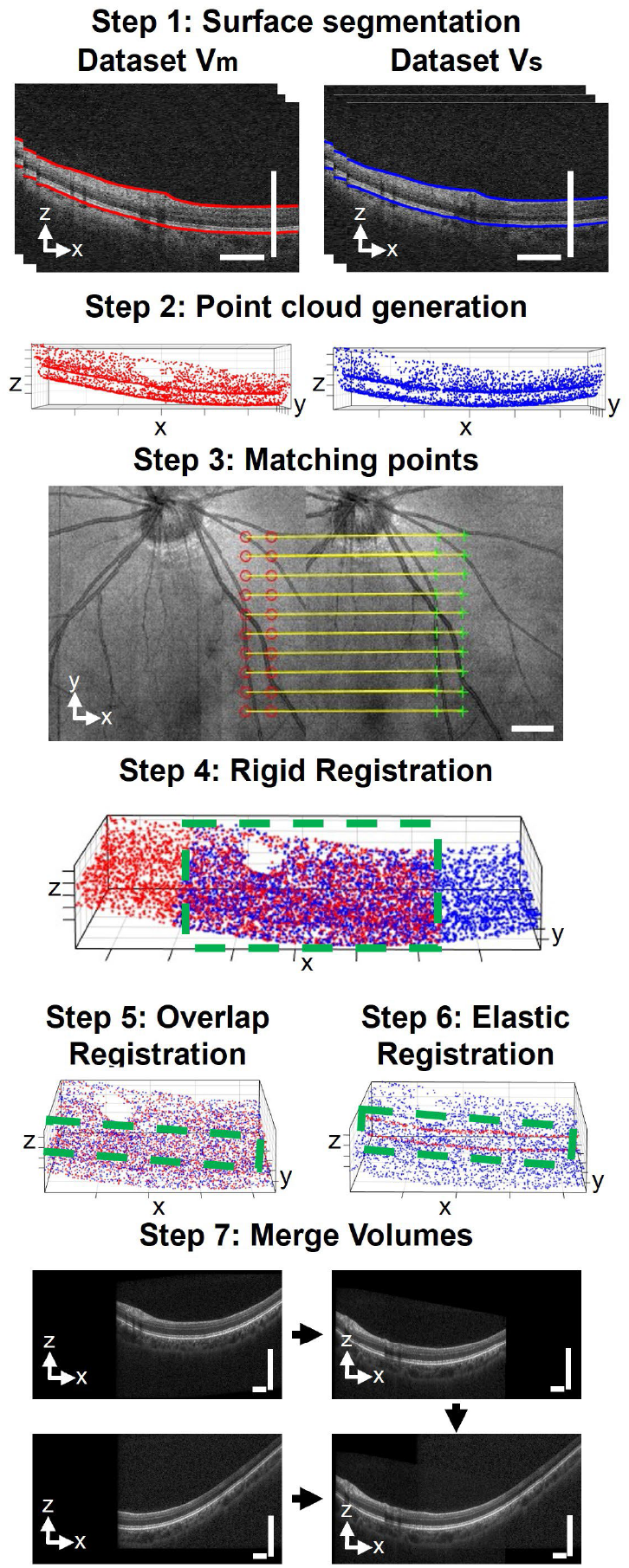
Flowchart for montaging adjacent volumes. Step 1: The ILM and RPE for each volume are segmented. Step 2: Point clouds are created from the spatial coordinates of the segmented ILM and RPE. Step 3: Matching landmark points between *V*_*m*_ and *V*_*s*_ are identified. Step 4: The matching landmark points and point clouds are used to generate an initial rigid transformation between *V*_*m*_ and *V*_*s*_. Step 5: The overlapping region between point clouds (green box in Step 4) is identified, and the transformation mapping the point clouds to each other in the overlapping region is computed to generate a final rigid transformation between *V*_*m*_ and *V*_*s*_. Step 6: Strip-based registration generates a deformable registration between *V*_*m*_ and *V*_*s*_. Step 7: The computed rigid and deformable registrations are applied to map overlapping volumes into the reference frame of *V*_*s*_. The scale bars for Steps 1, 3, 5, and 7 are 1 mm. The distance between ticks is 0.2 mm for the z axis and 1 mm for the x and y axes in Steps 2, 4, 5, and 6. Each point cloud was down-sampled by a factor of 20 for visualization.

### C. Step 1: Surface segmentation

We used a U-Net-based machine learning model to segment the ILM and RPE [34]. To create the model, we fine-tuned the existing OCT-Retinal Layer Segmenter model using 20 manually segmented B-scans. We trained the model on an NVIDIA GTX 3090 Ti GPU with 24GB of VRAM. During training, we minimized the categorical cross-entropy function

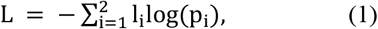

where *l*_*i*_ is the true label for sub-layer *i* and *p*_*i*_ is the predicted probability of sub-layer *i* from the model output. In our case, the two sub-layers comprised the retina between the ILM and RPE, as well as the background. We defined the ILM and RPE as the inner and outer layers of the segmented retina. **Step 1** in Fig. 1 shows the segmented ILM for two B-scans.

### A. Step 2: Point cloud generation

We generated a point cloud [35] of the 3D coordinates of the segmented ILM and RPE surfaces. We defined the point cloud as the spatial coordinates of the ILM and RPE boundaries across all B-scans. Alternatively, only one boundary, either ILM or the RPE, could be used when segmentation is challenging.

For *V*_*m*_, we defined *n* as the number of A-lines per B-scan, and *b* as the number of B-scans per OCT volume. We defined *x*_*i*_ as the x coordinate for each A-line within a B-scan and *y*_*j*_ as the y coordinate for each B-scan. We let *F*(*x*_*i*_, *y*_*j*_) and *G*(*x*_*i*_, *y*_*j*_) be the axial depths of the ILM and RPE at the lateral coordinate *x*_*i*_, *y*_*j*_. Using these definitions, the point cloud *M* for volume *V*_*m*_ is

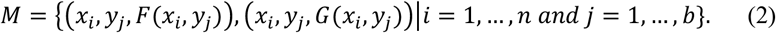

We defined the point cloud *S* for *V*_*s*_ identically.

### B. Step 3: Matching points

We determined the spatial coordinates of matching landmark points in the *en-face* projections of *V*_*s*_ and *V*_*m*_. **Step 3** of Fig. 1 shows matching landmark points in the *en-face* projections of *V*_*m*_ and *V*_*s*_. We generated a set of 2D grid points in the *en-face* image of *V*_*m*_, with the grid located either in the top, left, right, or bottom region of the *en-face* projection of *V*_*m*_, with the choice of region corresponding to where the *en-face* projections of *V*_*m*_ and *V*_*s*_ overlapped. For the example in **Step 3** of Fig. 1, the points located at the red circles correspond to the set of 2D grid points mentioned above. In this case, the grid points were located on the right because the right side of the *en-face* projection of *V*_*m*_ overlapped with the *en-face* projection of *V*_*s*_. The points in the green circles in Step 3 of Fig. 1 correspond to the matching points in the *en-face* projection of *V*_*s*_. We used normalized image screen coordinates [36] to describe the locations of the grid points, where the top-left point of the *en-face* projection corresponds to (0,0), the bottom-right to (1,1), the top-right to (1,0), and the bottom-left to (0,1). When the 2D grid points were in the left or right region, the grids consisted of two columns of ten points with a separation of (0.1, 0) between points in the x direction and (0, 0.075) between points in the y direction. The top left point of the left grid was located at (0.225, 0.2), and the top right point of the right grid was located at (0.775, 0.2). The descriptions of the grid points in the top and bottom regions are analogous, except that the x- and y-axes are swapped.

For each grid point, we took a 0.25 normalized width (25 percent of the *en-face* projection width) by 0.25 normalized height region from the *en-face* projection of *V*_*m*_ centered on the point. We computed the 2D normalized cross-correlation between this region [37] and the *en-face* projection of *V*_*s*_. We assigned the coordinate in the *en-face* projection of *V*_*s*_ with the maximum normalized cross-correlation as the matching point to the grid point in the *en-face* projection of *V*_*m*_.

### C. Step 4: Rigid registration

We obtained an initial estimate of the rigid transformation that maps the reference frame of *V*_*m*_ to that of *V*_*s*_ by computing the transformation that maps the spatial coordinates of common landmark points between *V*_*m*_ and *V*_*s*_. Rigid transformations preserve the spatial relationship between points, including translations, rotations, and reflections.

Given a total number of *p* matching landmark points, we defined the set of matching landmark points for *V*_*m*_ as

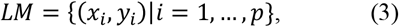

where (*x*_*i*_, *y*_*i*_) is the lateral position of the i^th^ landmark point.

We defined the set of 3D coordinates corresponding to these landmark points for *V*_*m*_ as

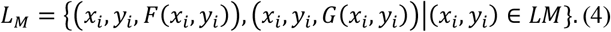

Similarly, we specified the matching feature points for *V*_*s*_ as *LS*, and the 3D coordinates of these points as *L*_*S*_. Next, we used the M-estimator sample consensus (MSAC) algorithm [38] to compute the transformation *T*_*init*_ mapping *L*_*M*_ to *L*_*S*_, as shown in Step 4 of Fig. 1.

### D. Step 5: Overlap registration

We refined the transformation *T*_*init*_ by applying the iterative closest point (ICP) algorithm [39] to the subset of *M* and *S* within the spatially overlapping region between *M* and *S* (green box in Step 4 of Fig. 1). To determine the spatially overlapping region, we found the polygon bounding *S* by computing the alpha shape associated with *S* [40]. We let *OM* be the set of points of *T*_*init*_ (*M*) with x and y coordinates enclosed within the polygon bounding *S*. Subsequently, we define the points in *M* within the overlapping region as

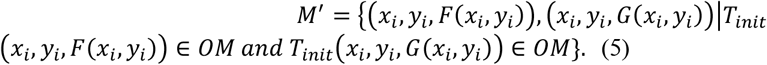

We determined *S’*, the region of *S* within the overlapping region, in an identical manner to *M’*. Finally, we applied the iterative closest point (ICP) algorithm to estimate the transformation T that maps *M’* to *S’*. Mathematically, the ICP algorithm minimizes an error function E

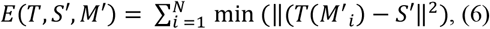

where the index *M’*_*i*_ is the i^th^ point of *M’* and ‖(*T*(*M*^′^_*i*_) − *S*′‖^2^ is the Euclidean norm of (*T*(*M*^′^_*i*_) − *S*′). Here, we let min stand for minimum, meaning that the closest point in *S*′ to *T*(*M*^′^) was used to calculate Euclidean distance. Step 5 of Fig. 1 shows T(*M’*) in blue and *S’* in red, with the two point clouds aligning closely in space.

### E. Step 6: Elastic registration

Due to eye motion, rigid transformations do not perfectly align the point clouds M and S. We adopted strip-based registration methods commonly used in adaptive optics OCT to compute a deformation vector field that best aligns the two volumes [41, 42]. Letting *a* and *b* be the upper and lower bounds for the y coordinates of the strip, we define the strip of *M* with y coordinates within these bounds as

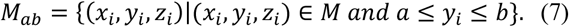

Next, we used the ICP algorithm to find the transformation *T*_*ab*_ mapping *M*_*ab*_ onto its associated region in *S* (Step 6 in Fig. 1). To prevent overfitting, we only used the subset *S*_*ab*_ of *S* consisting of the closest points in *S* to *T*(*M*_*ab*_). As with the calculation of the rigid transformation, the ICP algorithm minimized the error function E defined as

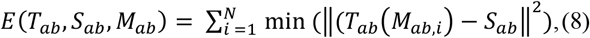

where *M*_*ab,i*_ is the i^th^ element of *M*_*ab*_.

Following the registration of each sub-strip of *M* onto *S*, we defined the transformation *T*_*s*_, consisting of a piecewise set of transformations aligning strips of M onto S. Letting *a* and *b* be the lower bounds for the first strip and *c* and *d* be the upper bounds for the last strip, we defined *T*_*s*_ as

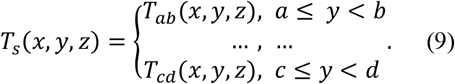

The displacement vector field *D* for each spatial coordinate (x, y, z) in the original frame is defined as

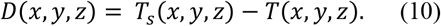

Conceptually, *D* is the displacement of the coordinates of *T*(*M*) necessary to align *T*(*M*) with the coordinates of *S*.

### F. Step 7: Merge volumes

Following the determination of *T* and *D*, we mapped the volumetric dataset of *V*_*s*_ and *V*_*m*_ to the common reference frame associated with *V*_*s*_ to form a merged volume *V*. As the merged volume *V* has the same reference frame as *V*_*s*_, we applied no transformations or deformations to *V*_*s*_. For volume *V*_*m*_, we updated its contribution to the intensity in V as

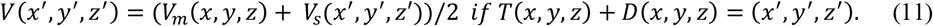

In other words, if coordinate (x, y, z) in *V*_*m*_ mapped to (x’, y’, z’) in *V*_*s*_ after applying the rigid and deformable transformation to *V*_*m*_, then the contribution of *V*_*m*_ to the merged volume *V* at the coordinate (x’, y’, z’) is *V*_*m*_(*x, y, z*).

In instances where more than two volumes contributed to the intensity *V*(*x, y, z*), we took the mean contribution from all the volumes.

Step 7 in Fig. 1 shows the process of merging the volumetric data. The upper left panel of step 7 shows a B-scan of *V*_*m*_ in its local reference frame. The rigid transformation *T* and displacement *D* are applied to map the *V*_*m*_ into the reference frame of *V*_*s*_. The lower left panel of Step 7 shows a B-scan of *V*_*s*_ in its local reference frame. The merged dataset of *V*_*m*_ and *V*_*s*_ combines the structural data of *V*_*m*_ and *V*_*s*_ in the reference frame of *V*_*s*_, with averaging of data in the overlapping region (bottom right panel of Step 7). In cases with three or more datasets, each volume is weighted equally in overlapping regions.

To generate *en-face* projections of the merged volume, we computed the mean of the 30 voxels with the highest intensity in the depth dimension at each lateral position. We chose not to use all voxels in the depth dimension for the *en-face* projection because each lateral position had a different number of voxels corresponding to background noise, depending on how many single volumes mapped to that lateral location.

### G. Assessing montaging error

Following 3D volumetric montaging, we generated a cumulative point cloud consisting of the union of every volume’s individual point cloud mapped into the common global coordinate. Suppose there exist point clouds *A* through *N*, with *T*_*A*_ being the rigid transformation for *A* into the global coordinate and *D*_*A*_ being the deformation vector field, then we define the cumulative point cloud CP as

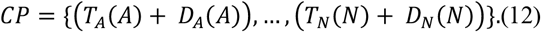

Next, we removed the point cloud corresponding to volume *V*_*A*_ from the cumulative point cloud. We determined the overlapping region between the point cloud *T*_*A*_(*A*) + *D*_*A*_(*A*) and the cumulative point cloud of the other volumes *CP* − *T*_*A*_(*A*) + *D*_*A*_(*A*) using the methodology described in **Step 5**. For every point of *T*_*A*_(*A*) + *D*_*A*_(*A*) in the overlapping region, we measured the distance between the point and the closest point in *CP* − *T*_*A*_(*A*) + *D*_*A*_(*A*). We defined the mean surface distance (MSD) as the mean distance between points in *T*_*A*_(*A*) + *D*_*A*_(*A*) and the nearest point in *CP* − *T*_*A*_(*A*) + *D*_*A*_(*A*). Additionally, we defined the median surface distance as the median distance between points in *T*_*A*_(*A*) + *D*_*A*_(*A*) and the nearest point in *CP* − *T*_*A*_(*A*) + *D*_*A*_(*A*). Similarly, we defined the 95% Hausdorff distance as the 95^th^ percentile of the distance between points in *T*_*A*_(*A*) + *D*_*A*_(*A*) and the nearest point in *CP* − *T*_*A*_(*A*) + *D*_*A*_(*A*). We repeated this process for every volume.

### H. Statistical analysis

All statistical comparisons between two groups were done with a paired T-test, and all comparisons between more than two groups were done with a one-way ANOVA. We reported all values as mean ± standard deviation.

## III. Results

### A. Montaging datasets from multiple institutions

We volumetrically montaged four subjects imaged at the HKU Eye Centre. Fig. 2a and 2b show the *en-face* projection for the montaged dataset from one subject, with the nine dotted squares highlighting the FOV for each original volume shown in Fig 2a. The FOV of the montaged volume in Fig. 2b was 18.2 mm × 17.4 mm and 3.2 mm deep. The FOV for a single volume in Fig. 2a was 9.3 mm × 9.3 mm and 1.9 mm deep.

**Fig 2.**
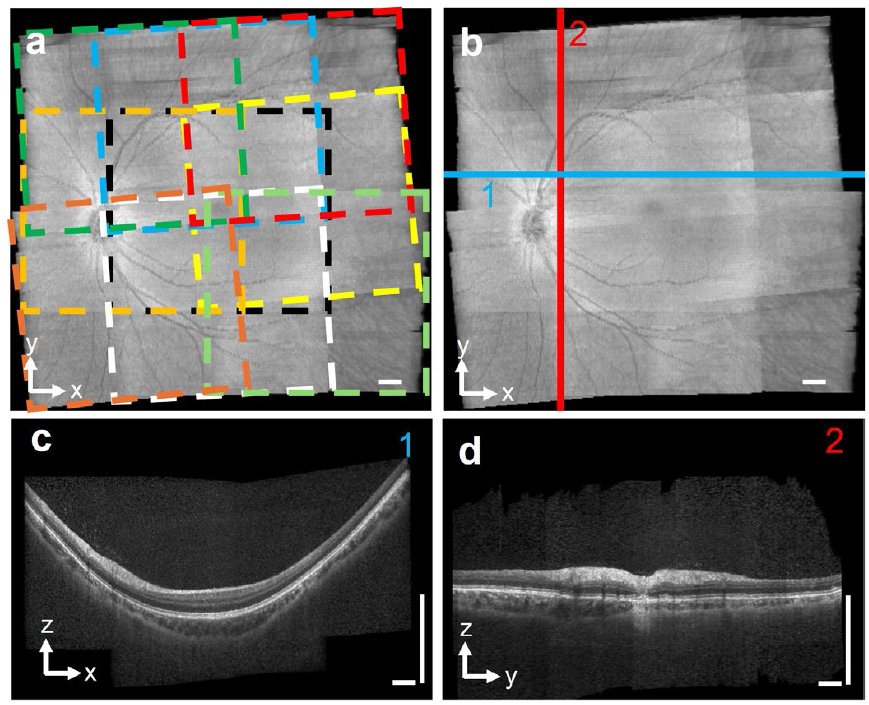
Montaged volume from HKU Eye Centre. a) *En-face* OCT (Spectralis2, Heidelberg Engineering) projection of montaged volume. The FOV consists of nine overlapping single-volume FOVs (dotted squares). b) *En-face* OCT without single-volume FOV overlayed. c) B-scan corresponding to the blue line in (b). d) B-scan corresponding to the red line in (b). Scale bars are 1 mm.

Fig. 2c shows a B-scan image from the montaged volume from the position highlighted by the blue line in Fig. 2b. Fig. 2d shows the B-scan image from the position highlighted by the red line in Fig. 2b. Both B-scan images show continuous retinal layers in the final montaged volume. We computed the average MSD between volumes as 38.3 ± 14.0 µm (n = 4 eyes) after rigid registration and 26.8 ± 13.3 µm after elastic registration.

We tested our montaging algorithms in eight eyes imaged at Wills Eye Hospital. Fig. 3a shows the montaged *en-face* projection, with dotted squares highlighting the FOV for each volume. The FOV for the montaged volume in Fig. 3 was 15.0 mm × 6.8 mm and 2.6 mm deep. The FOV for each volume was 6.0 mm × 6.0 mm and 2.0 mm deep.

**Fig 3.**
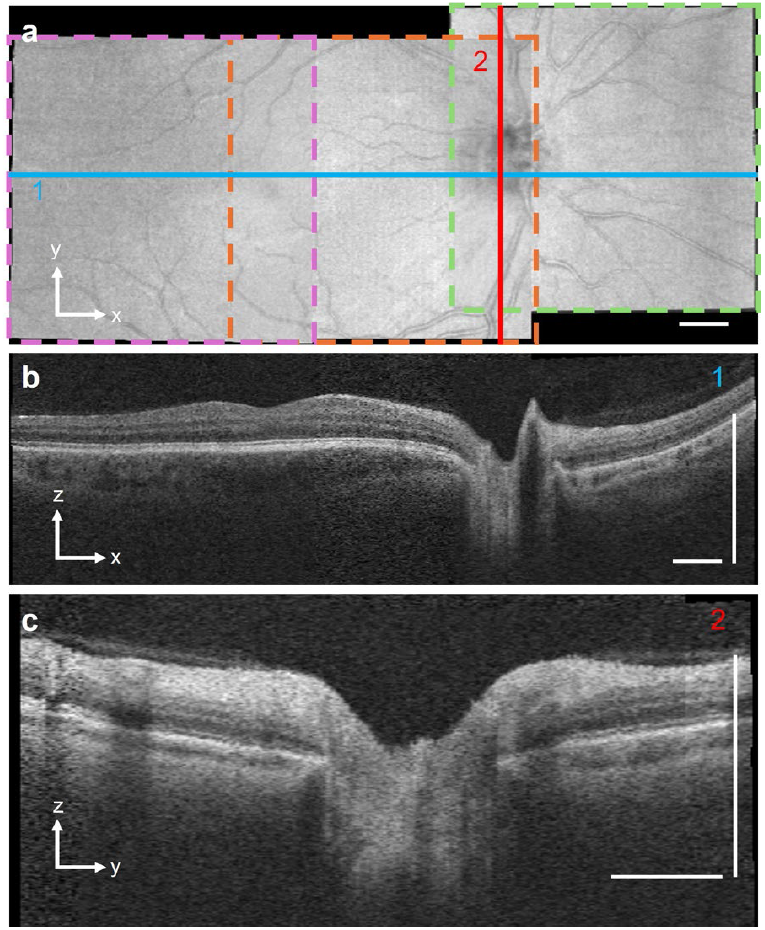
Montaged volume from Wills Eye Hospital. a) *En-face* OCT (Cirrus HD-OCT, Zeiss) projection of montaged volume, consisting of three single-volume FOV (dotted squares). (b) B-scan image from position highlighted by the blue line in panel a. (c) B-scan image from position highlighted by the red line in panel a. Scale bars are 1 mm.

Fig. 3b shows the B-scan image from the montaged volume from the position highlighted by the blue line in Fig. 3a. This B-scan integrates data from three volumes. Fig. 3c shows B-scan image from the position highlighted by the red line in Fig. 3a. Both B-scans contain data from multiple original volumes, none of them exhibit discontinuities at the boundary between volumes.

We montaged 18 eyes at Northwestern Memorial Hospital, with eight of the 26 eyes excluded due to significant motion artifacts. Fig. 4a and 4b show the montaged *en-face* projections, with dotted squares depicting the FOV for each volume in Fig. 4a. The FOV for the montaged volume in Fig. 4 was 17.2 mm × 12.2 mm and 3.5 mm deep. The FOV for each volume was 6.0 mm × 6.0 mm and 2.3 mm deep.

**Fig 4.**
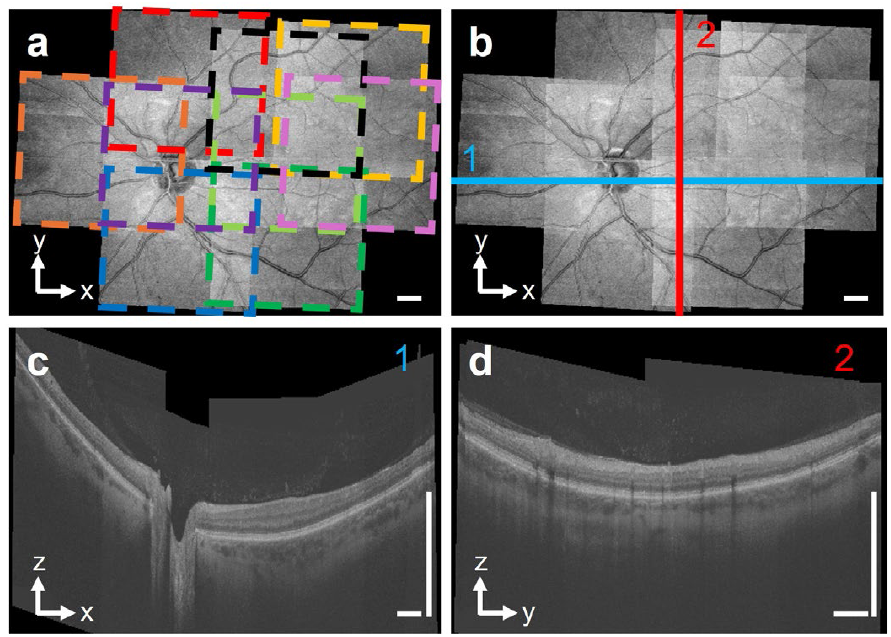
Montaged volume from Northwestern Memorial Hospital. (a) and (b) are *en-face* images (Maestro2, Topcon), consisting of nine overlapping volumes (FOV noted by dotted squares). (c) and (d) are B-scan images from the position highlighted by the blue and red lines in panel (b). Bar: 1 mm.

Fig. 4c shows the B-scan image of the montaged volume corresponding to the blue line in Fig. 4b, covering data from six volumes. Fig. 4d shows the B-scan image from the position highlighted by the red line in Fig. 4b, incorporating data from six volumes. Both B-scan image exhibit no discontinuities at the boundary between volumes.

### B. Large field of view montaged with 36 volumes

We acquired 36 volumes from a single eye at Wills Eye Hospital. Fig. 5a shows the *en-face* image of the montaged volume, with the dotted green squares highlighting FOVs of individual volumes. Fig. 5b shows the *en-face* image without the dotted boxes. The montaged volume covers 23.6 mm × 17.2 mm FOV and 4.8 mm deep.

**Fig 5.**
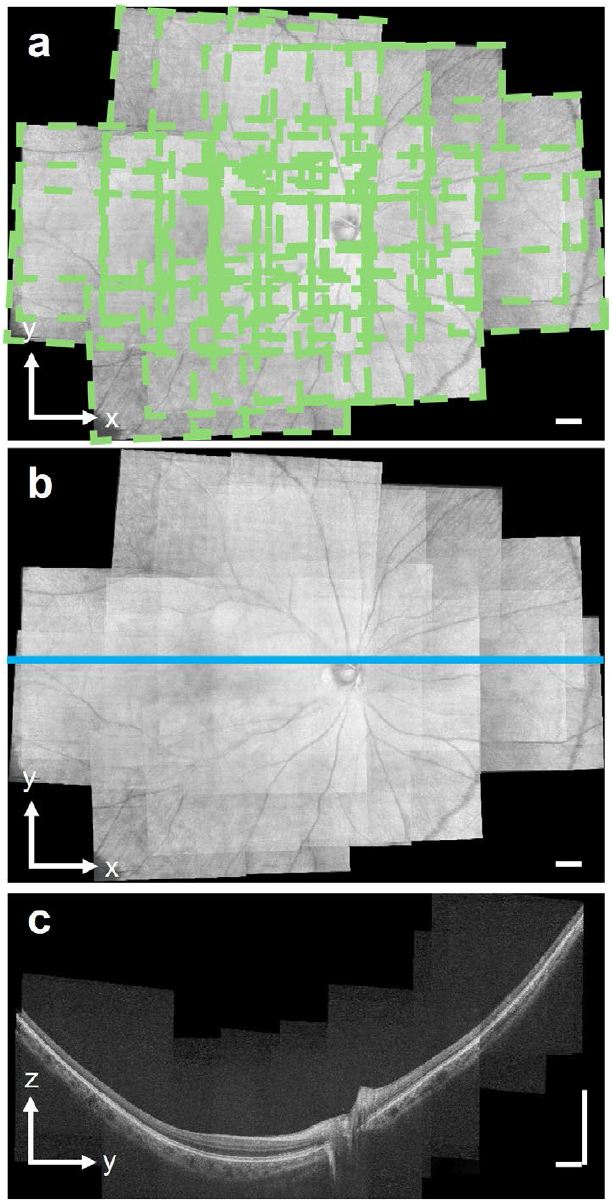
Volumetric montage of 36 volumes from a single eye. (a) *En-face* image of the montaged volume, consisting of 36 overlapping single volumes, with each FOV highlighted by the dotted squares. (b) *En-face* OCT without outlines of individual volumes highlighted. (c) B-scan image from the position highlighted by the blue line in panel b. Scale bar: 1 mm.

Fig. 5c shows the B-scan image of the montaged volume from the position highlighted by the blue line in Fig. 5b, incorporating data from 23 overlapping single volumes.

After rigid registration, we found that the MSD between volumes in this eye was 20.2 ± 18.3 µm (n = 36 individual volumes), the 95% Hausdorff distance was 68.2 ± 111.4 µm, and the median surface distance was 11.8 ± 5.5 µm. The maximum MSD was 82.8 µm, and the minimum MSD was 6.9 µm. After applying the elastic registration, we found that the MSD was 13.6 ± 12.2 µm, the 95% Hausdorff distance was 35.4 ± 54.4 µm, and the median surface distance was 9.5 ± 4.0 µm. The maximum MSD was 57.2 µm, and the minimum MSD was 6.4 µm. We observed that the MSD was less than the lateral pixel size of 30.15 µm but greater than the axial pixel size of 1.96 µm.

### C. Elastic registration decreases montaging errors

We compared the MSD after applying the rigid and the elastic transformations. Fig. 6a shows a B-scan image, with the ILM boundary highlighted in red. We show the point cloud corresponding to the ILM boundary below the B-scan. For the eyes (n = 26 eyes) acquired from Wills Eye Hospital and Northwestern Memorial Hospital, which had similar single-volume FOV, we performed volumetric montaging using the ILM point clouds. We found that the MSD was smaller (p = 0.02) after elastic registration (14.2 ± 9.1 µm) than after rigid registration (22.4 ± 22.8 µm).

**Fig 6.**
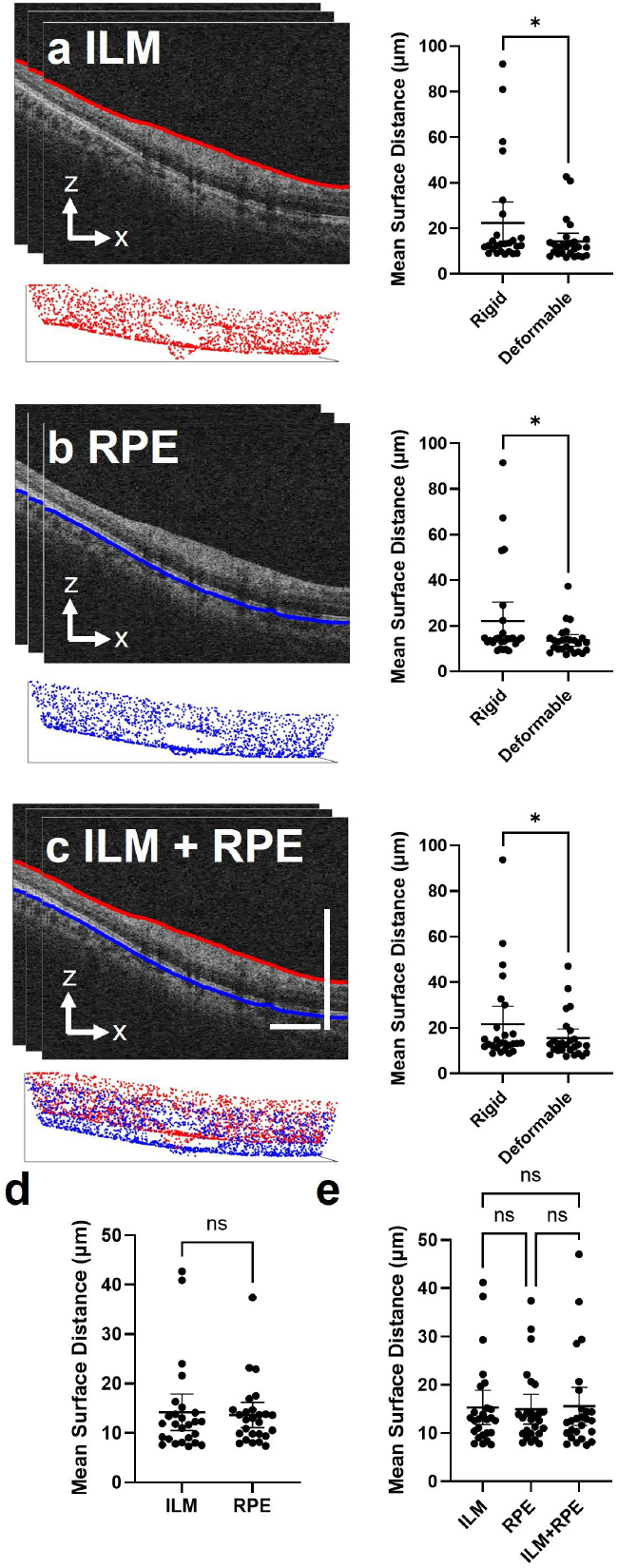
Comparison of point cloud generation methods. a) When point clouds were generated with the ILM border, MSD was smaller after elastic registration. b) When point clouds were generated with the RPE border, MSD was smaller after elastic registration. c) Multi-layer point clouds were generated with the ILM and RPE border, with MSD smaller after elastic registration. d) MSD for the RPE point cloud was comparable with the ILM point cloud. e) Comparable MSD for multi-layer point clouds when transformations were generated by single and multi-layer point clouds. Scale bars: 1 mm.

Fig. 6b illustrates a B-scan image with the RPE boundary highlighted by the blue curve. Below the B-scan, we show the point cloud corresponding to the RPE boundary. For the same set of eyes, we performed volumetric montaging using the RPE point clouds. We found that the MSD was smaller (p = 0.02) after elastic registration (13.6 ± 6.4 µm) than after rigid registration (22.1 ± 20.6 µm).

Fig. 6c shows a multi-layer point cloud generated using the ILM and RPE borders. The red surface depicts the topology of the ILM, and the blue surface depicts the topology of the RPE. We performed volumetric montaging using the multi-layer point clouds. We also found that the MSD was smaller (p = 0.04) after elastic registration (15.5 ± 9.7 µm) than after rigid registration (21.6 ± 19.4 µm).

### D. Comparison of point cloud generation methodology

We evaluated whether generating the point cloud using the ILM, the RPE, or both affected montaging errors. We performed volumetric montaging using point clouds generated from the ILM and measured the MSD of the cumulative ILM point cloud. We repeated the same procedure but used the RPE instead of the ILM. We found that the cumulative ILM point cloud MSD was 14.2 ± 9.1 µm (n = 26 eyes) and that the cumulative RPE point cloud MSD was 13.6 ± 6.4 µm (Fig. 6d), not statistically different than the ILM MSD (p = 0.49).

We further investigated the impact of the point cloud generation method on the MSD for the multi-layer cumulative point cloud. We found no statistically significant difference between the methodologies for generating point clouds (Fig. 6e). The MSD for the cumulative multi-layer point cloud was 15.3 ± 8.8 µm when the ILM point clouds were used to create the transformations, was 14.9 ± 7.7 µm when the RPE point clouds were used, and 15.5 ± 9.7 µm when the combined multi-layer point clouds were used.

## IV. Discussion

In this study, we presented a method for volumetrically montaging 3D OCT volumes acquired by three OCT devices (Spectralis2, Heidelberg Engineering; Cirrus HD-OCT, Zeiss; Maestro2, Topcon). We showed that 3D montaging of OCT volumes increased the FOV and axial depth without compromising high lateral sampling density. Most literature reports on OCT montaging and stitching were performed on 2D projection images. While scientists have learned much about the retina using stitched 2D projections, these methods fail to exploit the 3D nature of OCT fully. Other investigators have performed 3D registration *ex vivo*, primarily accounting for translation between volumes. We montaged 3D OCT retinal volumes from human subjects. We represented the ILM and RPE of each volume as point clouds and applied point cloud registration [35] to map the reference frames of each OCT volume into a common reference frame. By accounting for rotation, our method is robust to eye movement between acquired volumes, whose orientation to the incident OCT beam may differ substantially.

We tested our method using datasets acquired at three institutions using different OCT machines and acquisition protocols. For data from all institutions, we successfully expanded the FOV to more than 17 mm in the lateral directions. The increased FOV is comparable to that achievable with the latest ultra-widefield OCT, which has a FOV of around 20 mm in each lateral dimension [18, 19]. One benefit of 3D montaging is that it does not require new, expensive ultrawide OCT hardware. Since our methodology is not specific to any data-acquisition scheme, it can potentially be retroactively applied to historical data acquired from clinical OCT systems.

Two additional advantages of 3D montaging are denser sampling and an effective increase in axial imaging depth, which are usually constrained by either spectrometer design or the choice of a swept-source laser. For montaged datasets, the axial depth of the montaged volume was typically 1.5 to 2 times larger than that of a single OCT volume. We note that increasing axial depth helps minimize mirror artifacts [10] caused by insufficient imaging depth in OCT volumes, allowing individual regions without mirror artifacts to be montaged together. This increased imaging depth is particularly valuable in eyes with an elongated axial length, such as high myopia.

We quantified the montaging error by computing the MSD between the point clouds of the registered volumes. We found that the average MSD between point clouds after registration was less than the pixel size in the lateral dimension across all institutions, suggesting that the montaging algorithm maintains the accuracy of anatomical structure location up to the lateral spatial sampling distance of OCT. Additionally, nearly all A-lines in the montaged volume had well-distinguishable retinal layers. If any volumes were misaligned, we would have observed a misalignment of retinal layers for some regions of the montaged volume. Instead, we found that the structures of all volumes in the montaged volume aligned well.

We found that the MSD of the cumulative point cloud comprising the ILM and RPE was not statistically different, regardless of whether the ILM, RPE, or both layers were used to compute rigid and elastic transformations. This suggests that segmenting single layers of the eye is a reasonable alternative to using both layers to generate the point.

Additional applications for our volumetric montaging include compensating for motion artifacts within individual scans [43, 44] by treating regions of the volume acquired before and after motion as different volumes to be registered. Additionally, the volumetric registration method can potentially average volumes acquired at the exact location to increase signal quality [45]. Moreover, volumetric registration can merge volumes of the same region acquired at different angles, potentially improving the lateral resolution of OCT and bringing it closer to the axial resolution [46-49]. From a clinical perspective, we plan to conduct future studies to measure retinal curvature over a wider FOV, characterize the retinal topology in pathological myopia, and identify the 3D structure of peripheral lesions. Overall, our 3D montaging method will enable visualization of the retina’s 3D structure across a larger FOV and enhance understanding of how various disease states affect retinal structure.

## Acknowledgment

This research was supported by the NIH/NEI grants F30EY034033, U01EY033001, R01EY034740, R01EY038008, R01EY0130178, and R01EY030501. It was also supported by a seed funding from the Center for Engineering in Vision and Ophthalmology and an unrestricted departmental grant from Research to Prevent Blindness to Northwestern Ophthalmology.

